# Cessation of diel vertical migration by an inshore dinoflagellate bloom under prey deprivation

**DOI:** 10.64898/2026.07.16.738572

**Authors:** Serena Sung-Clarke, Nour Ayache, Wenguang Zhang, David K. Ralston, Evan Lechner, Zhaohui Aleck Wang, Juliette L. Smith, Collin Roesler, Susan Drapeau, Mengmeng Tong, Michael Brosnahan

## Abstract

Many dinoflagellates are mixotrophic and regulate their vertical position to navigate dynamic gradients in light, nutrients, and prey. Here, it is shown that the obligate kleptoplastidic mixotroph, *Dinophysis acuminata*, transitions from diel vertical migration to formation of a stationary, sub-surface thin layer in response to prolonged prey deprivation. An inshore bloom within a salt marsh kettle pond was recorded through continuous in-situ imaging, automated oxygen and fluorescence depth profiling, and targeted water chemistry measurements. During the bloom’s initial development, *D. acuminata* cells were photosynthetically active and divided vegetatively while vertically migrating. As photosynthesis and growth slowed, vertical migration ceased and cells formed a stable thin layer that promoted conditions for local acidification and nitrogen remineralization. Surface avoidance by the thin layer drove selective retention of cells within the relatively deep kettle hole. Together, these findings illustrate linkage of metabolic state and swimming behavior in *D. acuminata* and show how swimming behavior can drive development of toxic blooms within inshore systems. They also illustrate how *D. acuminata* and other eurytolerant bloom-forming species can exploit and shape physicochemical gradients associated with coastal eutrophication.

## Introduction

Phytoplankton thin layers— vertically narrow bands of high cell concentration—are subsurface hotspots of primary productivity and cell-cell interactions that promote genetic exchange and trophic transfers of energy and carbon within protist communities (Durham and Stocker 2012). Formation of these features is intricately intertwined with many species ability to vertically migrate across gradients of light and nutrient availability (Margalef 1983). Thin layers may also arise through interactions between cells’ swimming behavior and physical processes, including internal waves, wind stress, and tidal shear. These interactions can promote bloom development by concentrating cells within sharp physicochemical gradients or by enhancing aggregation (Largier 1993; Durham and Stocker 2012; Ralston et al. 2015).

The occurrence of thin layers is underestimated because their depth is often species-specific and typically only revealed when cell concentration is sufficient to cause a fluorescence anomaly (Schofield et al. 2008; Rines et al. 2010). Vertical migrations likewise have been masked by inadequate sampling approaches but are increasingly recognized through application of high-frequency, automated sampling methods (Garefelt et al. 2026). Vertical migration has been most intensively investigated in harmful algal blooms (HABs), particularly dinoflagellates (Schofield et al. 2008). Among these taxa, migratory behavior is adaptive across a wide range of conditions: cells may migrate over diel or longer periods to access deep nutrient reservoirs (Zheng et al. 2023) or alter swimming speed and preferred depths in response to changes in physiological state or life cycle stage (MacIntyre et al. 1997; Lewis et al. 2006; Brosnahan et al. 2017). The recognition that many vertically migrating protists are mixotrophic rather than purely phototrophic suggests additional drivers of migratory behavior including to enhance prey encounters and access of organic nutrient pools (Stoecker 1999; Stoecker et al. 2017).

This study examines thin layer formation and vertical migration in the HAB dinoflagellate *Dinophysis acuminata*, an obligate mixotroph that retains the plastids of its ciliate prey as kleptoplasts (Park et al. 2006; Hansen et al. 2016). *Dinophysis* species are globally important because they produce the diarrhetic shellfish poisoning (DSP) toxins okadaic acid and dinophysistoxins. These toxins accumulate in shellfish and can cause illness when contaminated shellfish are consumed (Reguera et al. 2014). Although all described species require light and ingestion of *Mesodinium* ciliates to grow in culture, blooms are frequently observed in situ in the absence of *Mesodinium* (Harred and Campbell 2014; Díaz et al. 2025). This includes Nauset Marsh (Cape Cod, MA, USA), a eutrophic estuary where prior work suggests bloom development can be driven by sexual reproduction rather than by prey ingestion and vegetative division (Sung-Clarke et al. 2026). The formation of *Dinophysis* blooms without *Mesodinium* may also be facilitated by physical aggregation; for example, stratification has been associated with the development of large-scale *Dinophysis* blooms and thin layers (Alves and Mafra 2018; Díaz et al. 2021). Whether blooms undergo diel vertical migration (DVM) remains unclear, however, as previous studies have reported both migrating and non-migrating populations (DVM; Villarino et al. 1995; Pizarro et al. 2008; Sjöqvist and Lindholm 2011).

Here, high frequency observations by a coupled continuous imaging-in-flow sensor and hydrographic profiler system were used to observe *D. acuminata* thin layer formation and migratory behavior through the development and decline of a bloom in situ. This approach resolves changes in DVM in response to prolonged prey deprivation, providing insight into how vertical migration may be altered to reflect changing cellular needs. In particular, cessation of DVM likely enhances survivorship during periods when cells can neither photosynthesize nor capture (or encounter) prey. Subsurface thin layer formation may also intensify and prolong DSP risk by enhancing cell retention within eutrophic coastal systems.

## Methods

### Study site

A *Dinophysis acuminata* bloom was recorded in 2024 within Salt Pond (Eastham, MA, USA), the northern terminus of the Nauset Marsh. Salt Pond is a relatively deep (maximum depth ∼10m) kettle pond that is connected to the greater Nauset Marsh and the Atlantic Ocean via a network of shallow (1-2m) tidal channels (Fig. 1). Salt Pond and other termini of the system are eutrophic due to substantial septic wastewater input via groundwater seepage (Giblin and Gaines 1990; Crusius et al. 2005; Colman and Masterson 2008).

**Figure 1.**
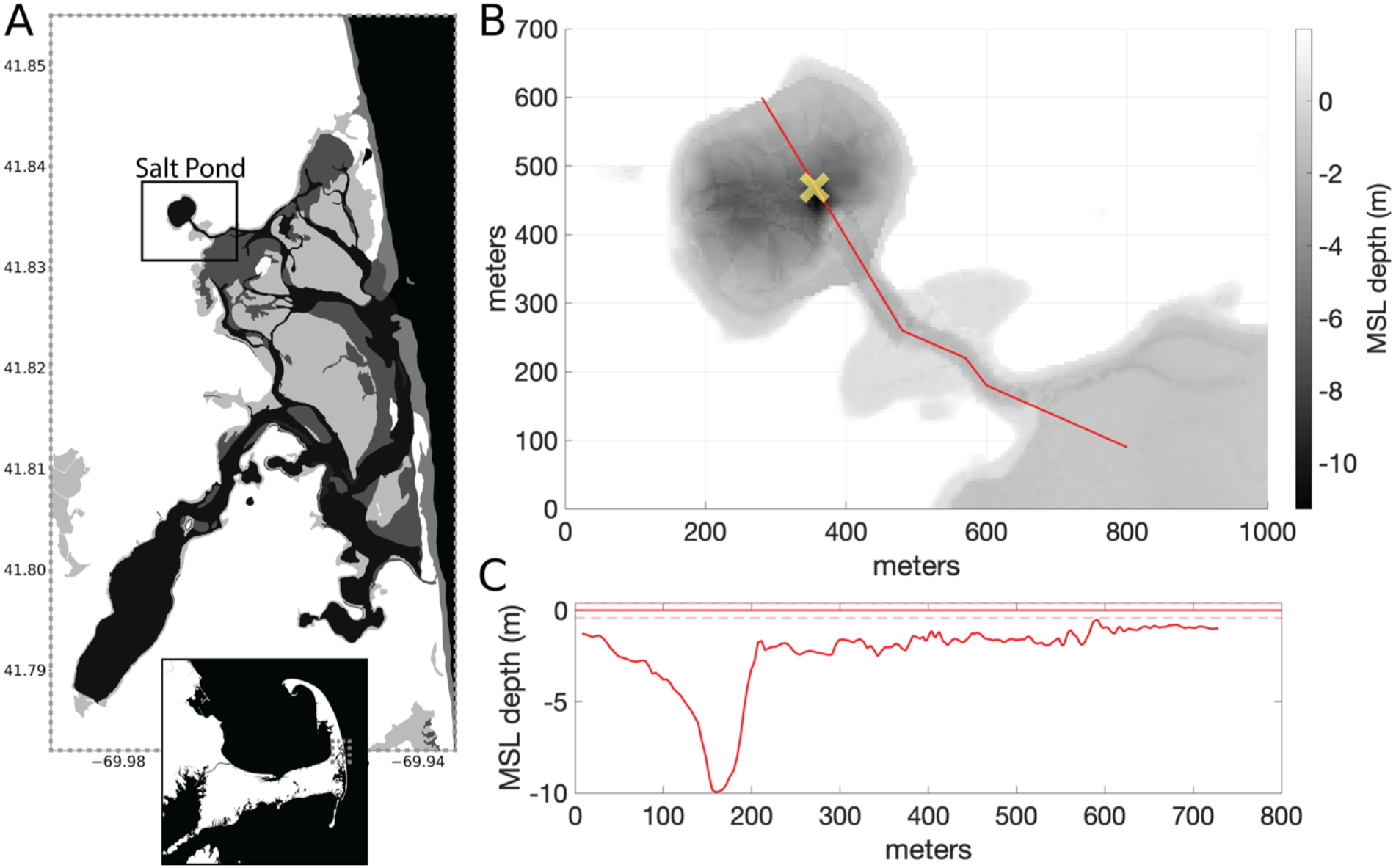
**(A)** Location and bathymetry of Salt Pond, the northwest terminal kettle pond in the Nauset Marsh, MA. **(B)** The bathymetry of Salt Pond. The “X” in the map indicates the location of the profiling observatory. Red line matches the transect described in (C). **(C)** depth across Salt Pond and into Salt Pond Bay.

Blooms of *D. acuminata* have been observed in Salt Pond nearly every year since 2015, when the first shellfish closure due to diarrhetic shellfish poisoning (DSP) was triggered upon observation of high *Dinophysis* concentrations by an Imaging FlowCytobot (IFCB) phytoplankton sensor (Ayache et al. 2023). The IFCB is a submersible imaging-in-flow cytometer that captures digital images of individual cells and other planktonic particles between 5 and 150 µm in length. Integrated antifouling features enable it to collect samples autonomously for months without recovery for maintenance, making it well-suited for observing *Dinophysis* and *Mesodinium* blooms (Olson and Sosik 2007; Campbell et al. 2010). Prior work has documented how surface avoidance by dinoflagellates interacts with the bathymetry and stratification at the site to drive selective retention of blooms (Anderson and Stolzenbach 1985; Ralston et al. 2015). Motivated by the association of this retention mechanism and HAB development, an observatory float is routinely deployed at the site for continuous bloom monitoring using an IFCB and automated fluorescence profiling (Fig 1., Brosnahan et al. 2017).

### In situ bloom recording through robotic profiling

In 2024, the Salt Pond observatory float consisted of an IFCB, an AML-6 sonde (AML Oceanographic, Victoria, BC, Canada), and a pair of winches that dynamically controlled both sonde profiling and depth of IFCB and water chemistry sample collections. The sonde was secured near the bottom of the IFCB so that it recorded relatively undisturbed water during downcasts of the combined IFCB-sonde unit. Additionally, the IFCB sample intake was extended so that it drew its samples immediately adjacent to the sonde’s main array of sensors. These sensors included conductivity and temperature (XCH2-CT-STD), pressure (XCH2-PRS-0050), dissolved oxygen (XCH2-DO2-JFE-FT-20), chlorophyll (XCH2-CHL-AB-TUR-06), and phycoerythrin fluorescence (XCH2-PHYE-TUR-20; AML Oceanographic, Victoria, BC, Canada). An additional photosynthetically active radiation (PAR) sensor (RSE-PAR-LIC-05-01; LI-COR, Lincoln, NE, USA) was attached to the auxiliary port of the AML-6 and secured above the IFCB to minimize shading from the IFCB and sonde. The sonde profile time series was quality controlled by discarding the measurements of individual sensors if they shifted abruptly, especially when such shifts were not accompanied by coincident shifts in other parameters.

Profiling of these sensors was controlled using a robot operating system (ROS) based middleware architecture called PhytO-ARM (https://github.com/WHOIGit/PhytO-ARM). The system was programmed so that a hydrographic profile from 1 to 6.5 meters was collected by dropping the IFCB-sonde package at a rate (∼2 cm s^-1^) prior collection of each IFCB sample. The IFCB was then positioned at the depth of maximum observed phycoerythrin fluorescence or a pre-programmed depth (1, 2.25, 3.5, 4.75, or 6 m), then triggered to collect and analyze a 5 mL seawater sample, so that both a sonde profile and IFCB sample were collected approximately every 25 minutes. IFCB image collection was configured to target red fluorescent particles (excitation/emission wavelengths: 635/680nm) so that records were not overwhelmed by high loads of non-fluorescent detritus. Samples were excluded from analyses if aggregate particle positions within its camera field of view indicated non-quantitative recording, or if its imaging rate dropped significantly, indicating hardware malfunction.

Tidal amplitude and additional temperature data were measured using a HOBO pressure logger (LI-COR, Lincoln, NE, USA) affixed to the bottom of Salt Pond near its northern shore (∼ 2m mean sea level). To derive relative tidal height, mean tidal height recorded while the logger was deployed (September 2023 – September 2024) was subtracted from each tidal height measurement. Wind speed and direction throughout the duration of the bloom were derived from the Provincetown Municipal Airport (42.07°N, 70.22°W), ∼34 km NNW of Salt Pond.

### Nutrient, carbonate, and pH analysis

In addition to continuous observations through the coupled IFCB-sonde system, samples for measurement of pH, dissolved inorganic carbon (DIC), and dissolved inorganic nutrients were collected every 5-14 days through the duration of the *Dinophysis* bloom (June 7 – July 24). On each sampling date, a peristaltic pump with inline 0.45µm and 0.2µm capsule filters in sequence (Farrwest Environmental Supply, Texas, USA) was connected to plastic tubing on the platform’s secondary winch. On June 7 and 12, samples were collected at the depth of the phycoerythrin maximum. Samples were collected only during ebb tide on June 7 and during both flood and ebb tides on June 12. From June 20 onward, sampling was conducted at slack high tide and from five depths, in increments of 0.25 m from 0.5 m below to 0.5 m above the position of the phycoerythrin maximum.

Bottle sample collection and measurements for DIC and pH followed the best practice of seawater CO2 measurements (Dickson et al. 2007). Filtered samples for measurement of DIC and pH were collected in separate 100 mL borosilicate glass bottles, poisoned with saturated mercuric chloride (to 0.04% v/v), then sealed with an air-tight screw top cap in the field. DIC measurements were made with a AS-C6L DIC multi-sample analyzer (Apollo SciTech Inc., Delaware, USA). Measurements of pH were made with an Agilent 8453 following standard spectrophotometric procedures (Supporting Information Text S1). Overall measurement uncertainties for DIC and pH were ±0.1% and ±0.006 pH units, respectively. Water for nutrient analyses was collected in duplicate acid-washed plastic bottles and stored on ice for transport back to the laboratory (<4 h), then transferred to a −20°C freezer for storage until analysis at Bowdoin College for ammonium, total oxidized nitrogen (nitrate + nitrite), and phosphate using a SmartChem autoanalyzer with methods adapted from Westco Unity SmartChem Methods, with uncertainties for nitrogen and phosphorus measurements ±0.1µM and ±0.01µM, respectively (Supporting Information Text S2).

### IFCB image analysis

IFCB images were classified through a two-step process. Images were first sorted taxonomically with an inception_v3 convolutional neural net model (Szegedy et al., 2015) that examined 93 taxonomic groups (44 species level, 45 genus, 2 order, 2 class) and 25 additional generic and non-taxonomic categories (e.g., pennate, detritus, fiber). Hourly samples at least every three days when available (∼24% of IFCB samples collected during the bloom) were then reviewed and corrected for final classification of *Dinophysis acuminata* and *Mesodinium* and identification of *Dinophysis* division and mating cells. These revisions were conducted using a web-based image labeling tool (https://ifcb-annotate.whoi.edu). Changes in mean cell appearance were evaluated using cell biovolume and average pixel grayscale intensity of *Dinophysis* images through calculation of v2 image blobs and features (https://github.com/hsosik/ifcb-analysis). In particular, changes in average pixel grayscale intensity were examined based on prior work that linked this parameter to recency of *Dinophysis* feeding (Ladds et al. 2024). Biovolumes were computed in voxels (Moberg and Sosik 2012), then converted to cubic microns using a conversion factor of 2.8 pixels per micron based on prior calibration for similar sized dinoflagellates (Brosnahan et al. 2015).

### Analysis of phycoerythrin fluorescence profiles

A log-log linear regression was used to describe the relationship between IFCB-derived *Dinophysis* cell concentration and simultaneous measurements of phycoerythrin fluorescence for all annotated IFCB samples (Eq. 1). This relationship, where β_0_ is the y-intercept and β_1_ is the slope, was then applied to phycoerythrin fluorescence values (*P*) at each timepoint (*i*) to estimate cell concentration within 20-cm depth bins (*j*) through the full depth of each fluorescence profile. To estimate standing stock of cells in Salt Pond, it was assumed that cell concentration at a given depth was horizontally uniform across the pond. For each timepoint *i*, the standing stock at that time, *N_i_*, was estimated by multiplying estimated concentration within a depth bin, *C_i,j_*, by the pond’s area at that depth, *A_j_*, and the thickness of each depth bin, Δ*z_j_*, then summing across all depths (Eq. 2). The area of Salt Pond as a function of depth, *A_j_*, was obtained from previously published bathymetric data (Ralston et al. 2015).

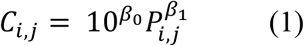

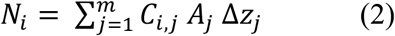

Phycoerythrin fluorescence values were averaged into 1-cm bins to increase measurement precision, then smoothed using a 5-bin rolling mean. The vertical position of the thin layer was evaluated as the maximum of this smoothed phycoerythin fluorescence profile. The vertical compactness of the thin layer within each profile was quantified to 5-cm resolution using a full width at half maximum (FWHM) approach, calculated from the depths above and below the fluorescence maximum where the smoothed signal was half its maximum value.

### Growth, accumulation, and loss rate derivations

For every day with IFCB annotations covering >18 hours, a *Dinophysis* growth rate was calculated. The intrinsic growth rate (*μ*) is the sum of its asexual (vegetative) division and sexual reproduction rates, *μ_d_* and *μ_s_*, respectively (Eq. 4), which were each calculated from observed frequencies (*f*) of dividing and mating cells and the duration of these processes (*t*) over *n* samples (Eq. 5). Division and mating durations were taken from Sung-Clarke et al. (2026) based on similarity of observed temperatures to the 15°C set point used in their experiments.

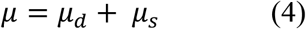

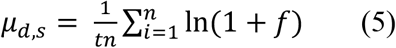

This intrinsic growth rate was then applied to phycoerythrin-derived cell abundance, *N_i_,* to project predicted change in abundance in the absence of other gain and loss processes. The relative contribution of asexual and sexual reproduction to cell growth was estimated as described in Sung-Clarke et al. (2026).

Intrinsic growth rates were compared to accumulation rates, *α*, from the same days, which were calculated from changes in standing stock estimates, where *N_t_* is the daily mean standing stock on day *t* and *N_t_*_+Δ*t*_ is the daily mean standing stock estimate at the subsequent day for which growth rates were available.

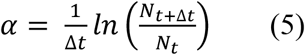

When biological growth dominates population gains relative to advective gains, the apparent loss rate (*l*) of the population is the difference between intrinsic growth (*μ*) and observed accumulation (*α*) rate.

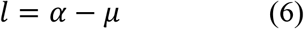

Lower-bound estimates of *Dinophysis* residence time during the bloom were estimated by fitting an exponential decay curve to changes in standing stock when intrinsic growth was near-zero (Supporting Information Text S3).

## Results

### An intense, dominant, and sub-surface Dinophysis bloom

From June 7 – July 24, 1,812 CTD profiles were collected. All sensors functioned continually throughout this period with the exception of the dissolved oxygen sensor, which ceased collecting reliable data after July 1. More than 13 million images were generated from 1460 IFCB samples collected while CTD profiling was active. The overwhelming majority of these samples (1182) were sampled at the phycoerythrin maximum depth, and the rest were sampled at preset depths: 67 at 1 m, 57 at 2.25 m, 55 at 3.5 m, 50 at 4.75 m, and 49 at 6 m.

A total of 286 IFCB samples were manually annotated. From this set, a total of 624,838 *Dinophysis acuminata* cells (including 2,509 dividing and 982 mating cells) and 143 *Mesodinium* cells were identified. The average pixel grayscale intensity distribution of *Dinophysis* images was highly consistent throughout (Supporting Information Fig. 1A). IFCB-derived estimates of *Dinophysis* concentration were strongly correlated with concurrently measured phycoerythrin fluorescence (R^2^ = 0.75, p << 0.01, Fig. 2B). Among the manually annotated IFCB samples, *Dinophysis* accounted for over half of the total biovolume recorded, including >90% in samples collected near the bloom’s peak (Fig. 2A).

**Figure 2.**
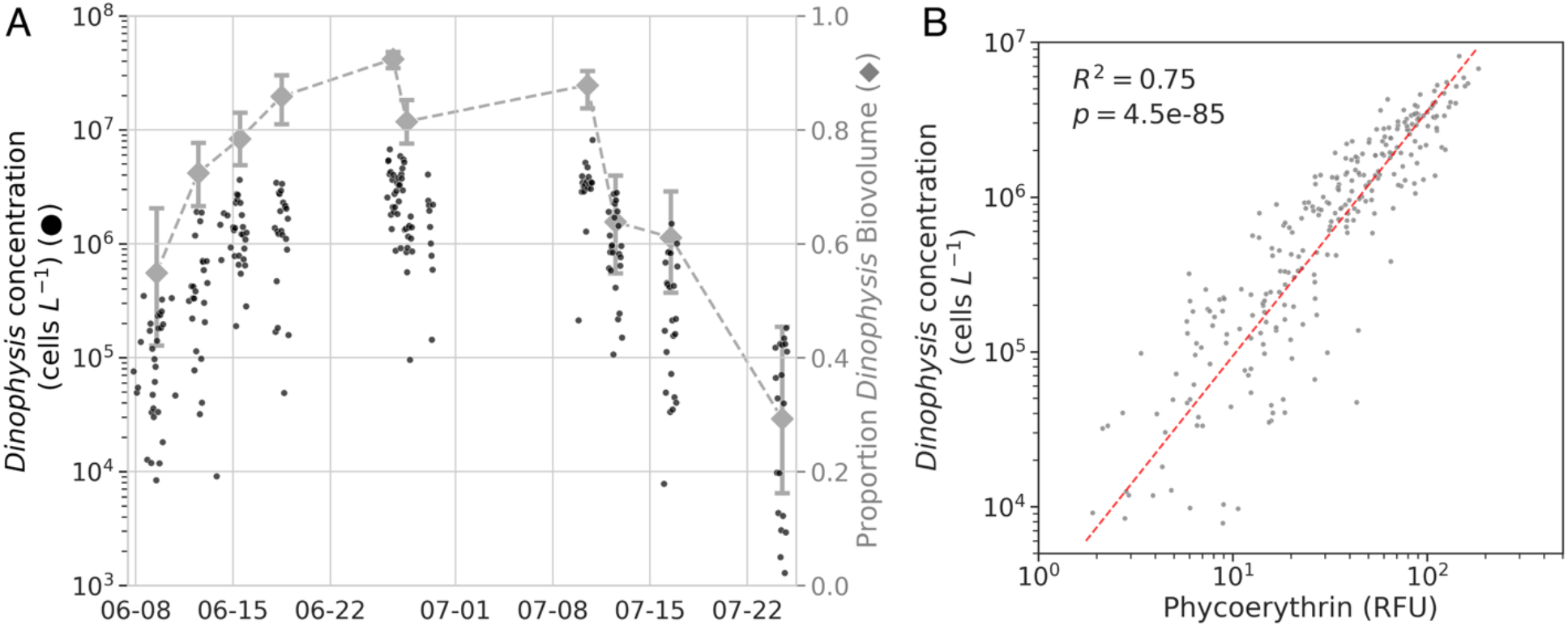
**(A)** IFCB-measured concentration of *Dinophysis acuminata* throughout the 2024 bloom, and the proportion of all IFCB image biovolume attributable to *D. acuminata* on each fully annotated day. **(B)** Correlation between concurrent IFCB-measured *D. acuminata* cell concentration and phycoerythrin fluorescence.

Phycoerythin fluorescence profiles revealed that *Dinophysis* were consistently localized within a sub-surface thin layer between 3 and 5 m deep and typically <1 m thick (Fig. 3B). Mean *Dinophysis* abundance estimated from depth-integrated fluorescence followed a trajectory very similar to concentrations calculated from IFCB images at the phycoerythrin maximum (Fig. 2A, Fig. 4A). Most new cell production during the bloom (∼80%) was attributed to vegetative division rather than sexual reproduction.

**Figure 3.**
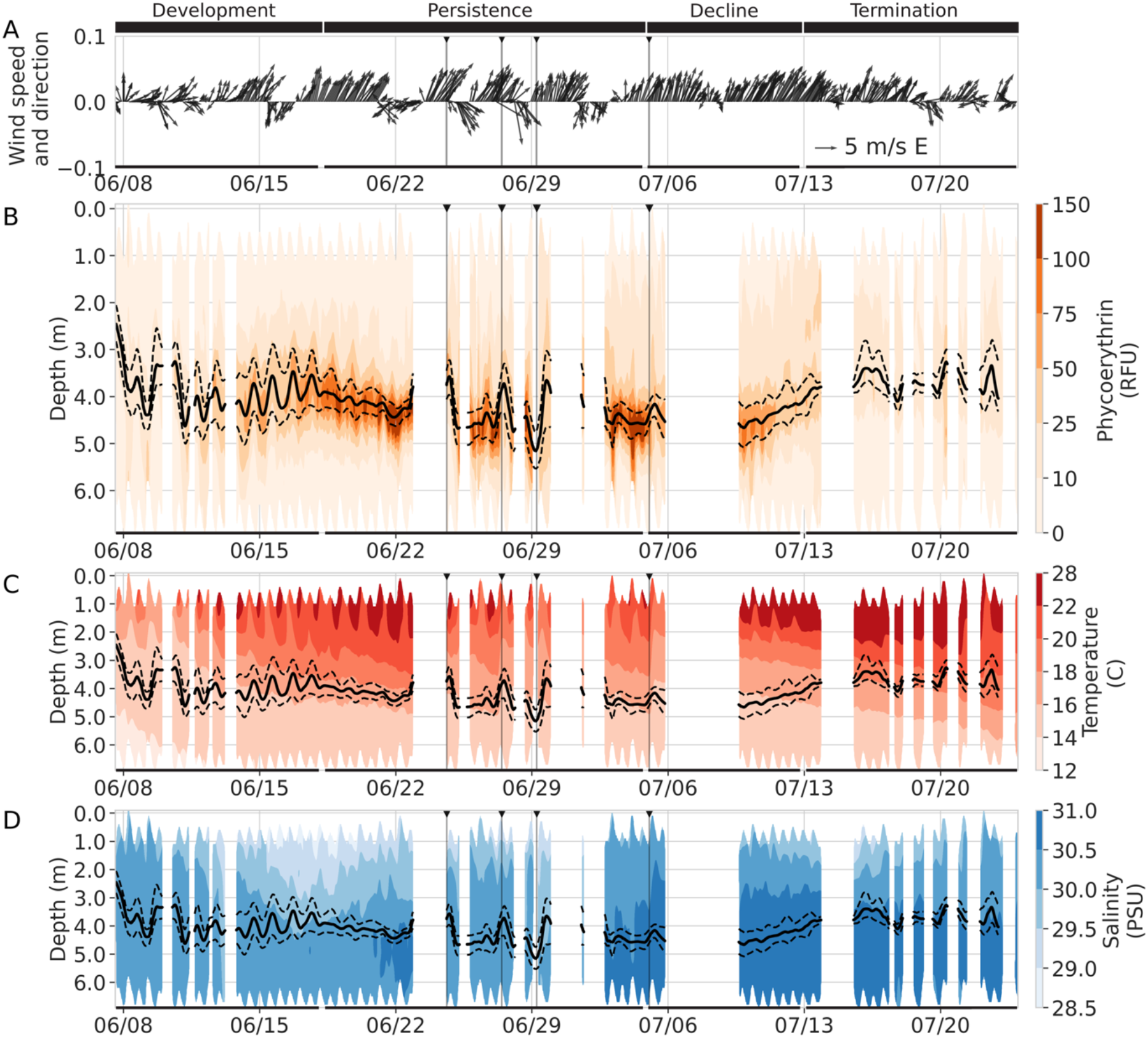
Physical conditions in Salt Pond during the 2024 *Dinophysis* bloom. **(A)** Wind speed and direction, reference arrow depicts 5m/s wind toward the east. **(B)** Phycoerythrin depth profiles (in relative fluorescence units, RFU). Gray solid line is the smoothed peak phycoerythrin depth and dashed lines represent the upper and lower bounds for the full-width at half-maximum from the peak. **(C)** Temperature and **(D)** salinity profiles throughout the bloom, with the black solid and dashed lines representing the smoothed depth of the phycoerythrin maximum and its upper and lower depth bounds. Depth is the depth below the mean sea level surface height. Vertical lines highlight four thin layer disruptions during the peak of the bloom.

**Figure 4.**
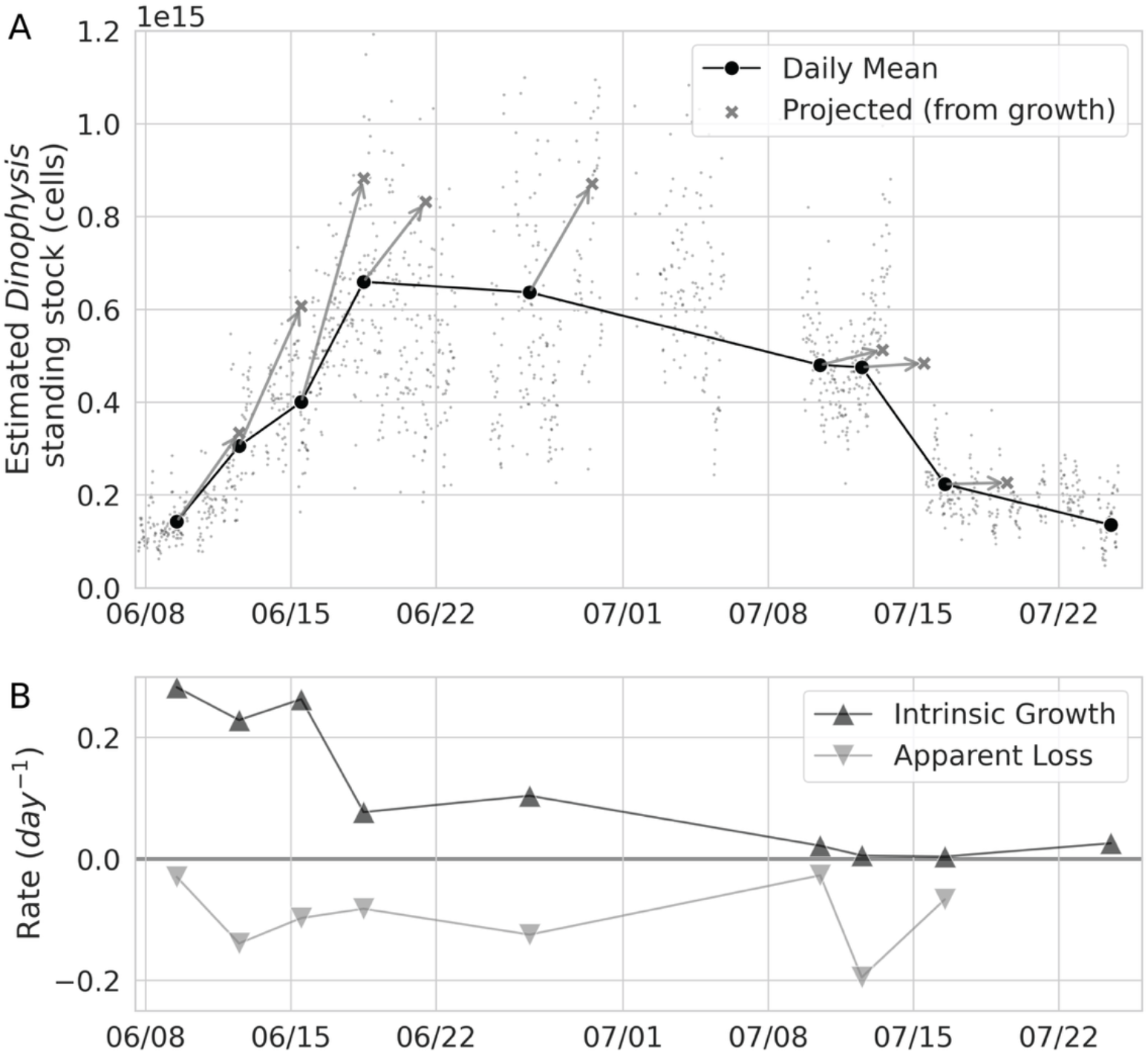
**(A)** Estimated *Dinophysis* standing stock in Salt Pond, derived from phycoerythrin profiles, with 3-day projected abundances from daily means based on cell division and mating frequency in IFCB imagery. **(B)** Growth rates derived from cell division and mating frequency and the loss rate defined as the difference between growth rate and the rate of cell accumulation.

IFCB-derived growth rates and phycoerythrin fluorescence-derived standing stock estimates were used to define four distinct bloom phases: development, persistence, decline, and termination. During the development phase (June 7-17), *Dinophysis* were actively growing (0.23 – 0.28 day^-1^) and the standing stock increased over five-fold. During the persistence phase (June 18 – July 4), growth rates were lower (∼0.08 – 0.11 day^-1^), thin layer cell concentration was consistently >1M cells L^-1^, and standing stock size was stable (Fig. 2A). During the decline phase (July 5-12), growth rates were near-zero (<0.03 day^-1^), and standing stock declined modestly (∼20%). In the termination phase (July 13-23), growth rates were similar to that in the decline phase (<0.03 day^-1^), but standing stock declined rapidly (Fig. 4A-B). Each of these phases was characterized by a distinct combination of physical conditions, thin layer characteristics, and chemical conditions (dissolved oxygen, inorganic nutrient concentrations, pH, and DIC).

### Vertical migration associated with phototrophic growth (June 7 – 17)

Growth during the development phase of the bloom was dominated by vegetative division, accounting for ∼85-90% of new cells). *Dinophysis* underwent modest vertical migrations, swimming ∼1m upward at dawn and ∼1m down at dusk (Fig. 3B). During the day, at their shallower depth, cells were exposed to PAR levels of ∼10-60 µmol photons m^-2^s^-1^, nearly 10-fold higher than at the depth of their ∼1m deeper nighttime position (light attenuation coefficient, *K_d_*, ∼ 2 m^-1^, Fig. 5A). Photosynthetic activity of vertically migrating *Dinophysis* was apparent from daily enrichment of dissolved oxygen in the thin layer, where *Dinophysis* was dominant (Fig. 5B-C).

**Figure 5.**
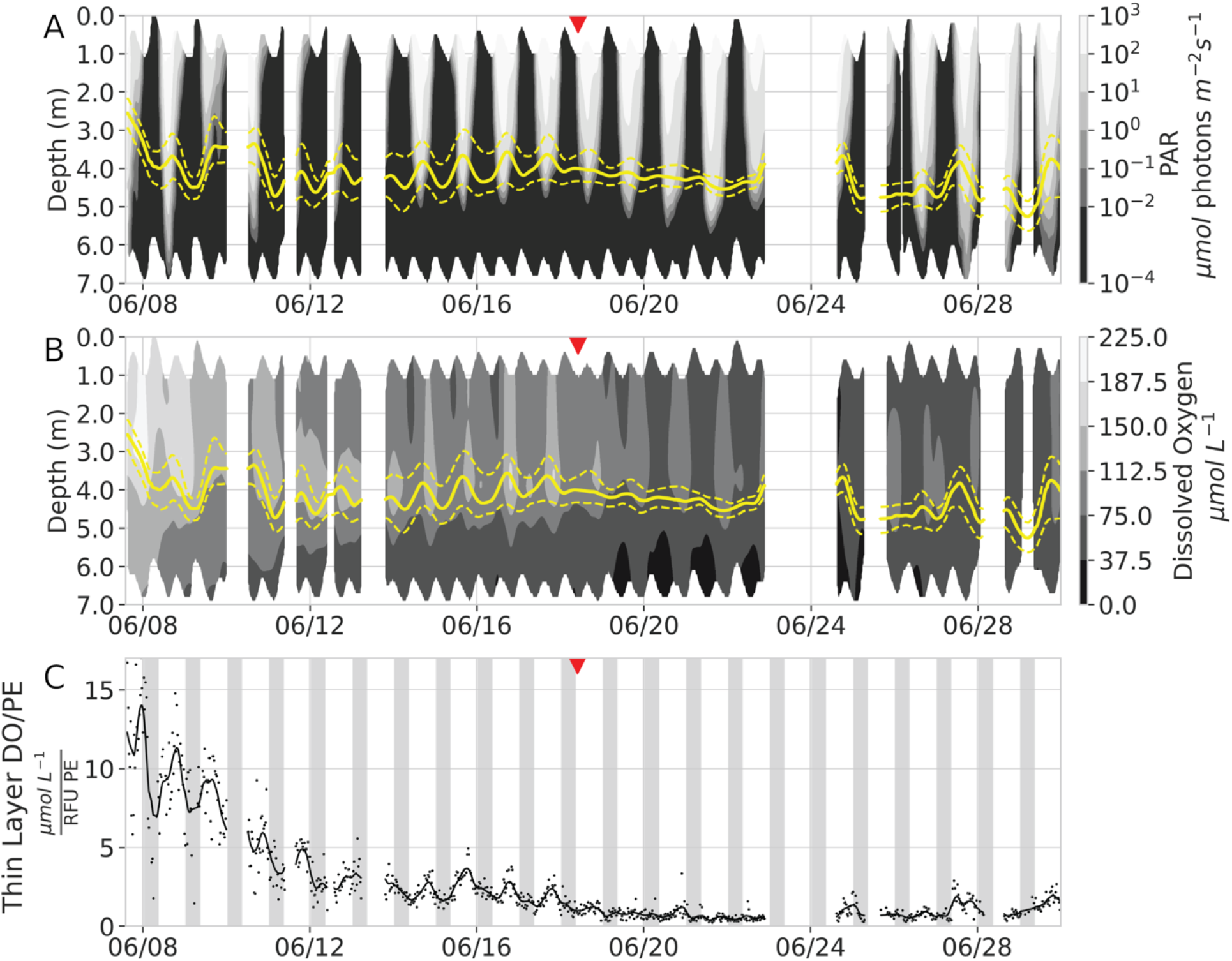
**(A)** Photosynthetically active radiation (PAR) and **(B)** dissolved oxygen profiles during bloom development and photosynthetic decline. Solid and dashed yellow lines represent the peak and upper/lower bounds of the *Dinophysis* thin layer. **(C)** Mean dissolved oxygen per mean relative fluorescence unit of phycoerythrin in the thin layer. Grey background bars represent nighttime periods from approximate sunset to sunrise. The small red arrow indicates the shift from the development to the persistence phase of the bloom, where diel vertical migration ceased.

The thin layer was ∼1.1m thick (FWHM) and generally resided in colder and saltier water below the pycnocline (14-18°C, salinity 29.5-30.5) (Fig. 3C-D, Fig. 6, Supporting Information Fig. 2B). Around the phycoerythrin maximum on June 7 and June 12, ammonium concentration was <1µM, phosphate <5µM, and nitrate/nitrite <0.8µM (Fig. 7). On June 7, thin layer pH was ∼7.6, and DIC was ∼1990-2020 µmol kg^-1^. On June 12, thin layer pH was 7.7-7.4 (top to bottom, Fig. 7) and DIC was 1950-2150 µmol kg^-1^ (top to bottom; Supporting Information Fig. 3).

**Figure 6.**
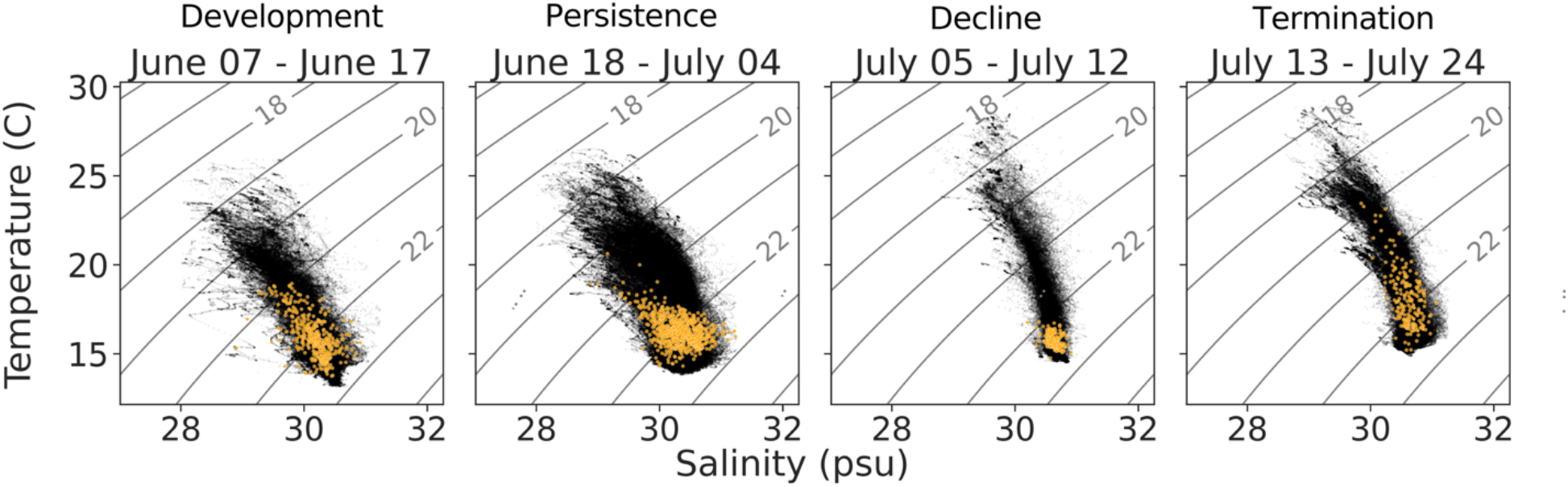
Temperature-salinity diagrams of the vertical profiles throughout each phase of the bloom, where black dots represent all measurements throughout the depth of the pond, and the orange dots are the temperature and salinity at the phycoerythrin maximum depth.

**Figure 7.**
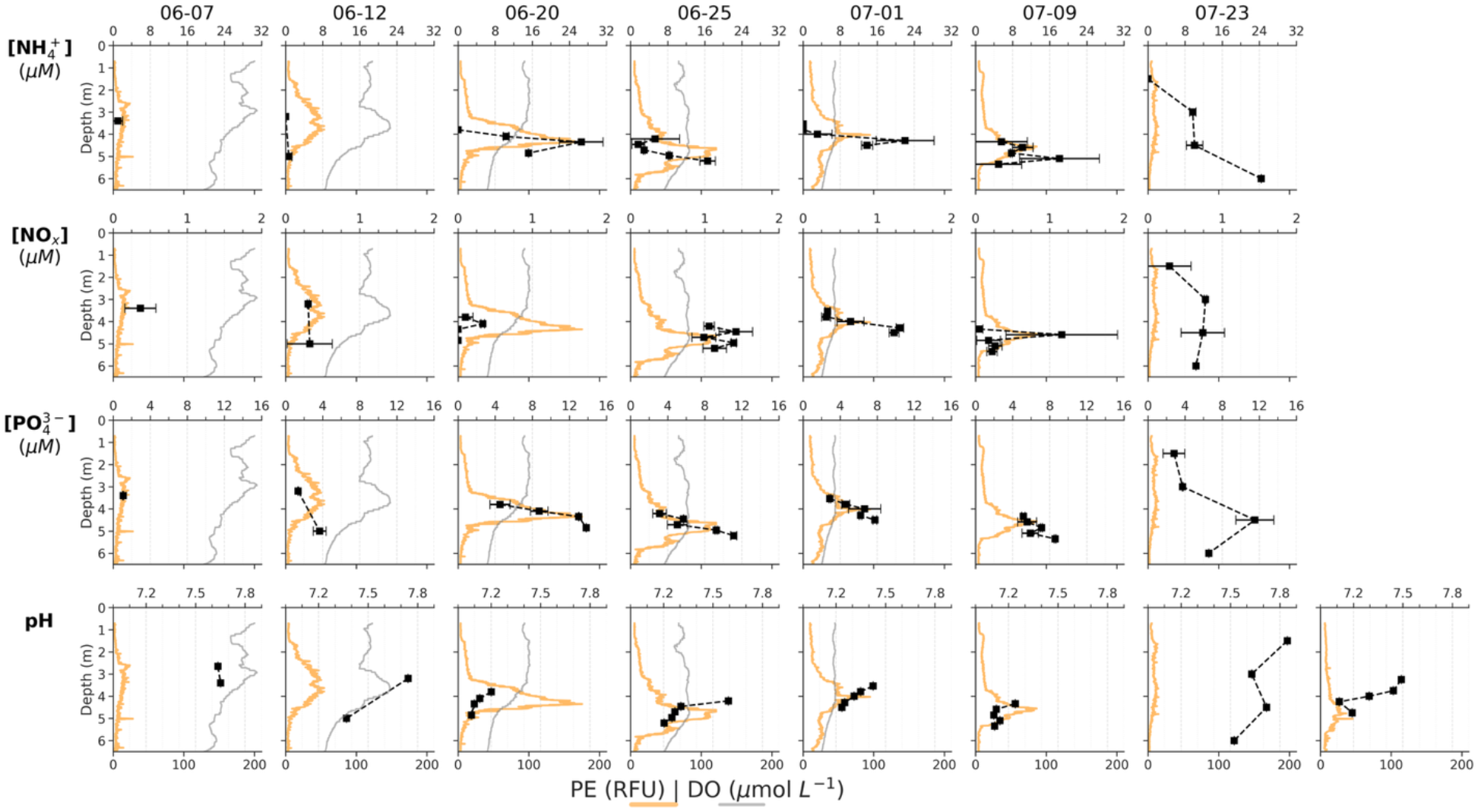
Ammonium (NH_4_^+^, top row), nitrate/nitrite (NO_x_, second row), phosphate (PO_4_^3-^, third row) concentration, and pH (fourth row) around the thin layer compared to phycoerythrin (PE) and dissolved oxygen (DO) profiles across the bloom. Top x-axes represent nutrient concentration or pH values; bottom x-axis values are phycoerythrin (RFU) and DO (μmol L^-1^) values.

### Transition to formation of a stable sub-pycnocline thin layer (June 18 – July 4)

Growth continued but was more modest and standing stock estimates stabilized during this period, indicating balance between growth and loss processes (Fig. 4B). Estimated loss rates were similar to those during the development phase (Fig. 4B). Sexual reproduction became a larger contributor to new cell production, but vegetative reproduction still dominated, accounting for ∼70-85% of new cells. *Dinophysis* cells did not vertically migrate; instead, they maintained a stable thin layer between 4 and 5 m below mean sea level depth (PAR ∼5-65 µmol photons m^-2^s^-1^). The thin layer also narrowed (mean FWHM 66cm; Fig. 3).

As vertical migration stopped, daytime oxygen enrichment became negligible despite little change in cell abundance (Fig. 5B-C). This decline in oxygen indicated a shift toward greater respiratory demand within the thin layer relative to photosynthetic oxygen output (Fig. 5B-C). Consistent with this shift, pH within the thin layer was low (7.1-7.3, Fig. 7) and a local maximum in ammonium concentration was observed at or just below it (15-30µM; Fig. 7). Nitrate+nitrite concentrations, although consistently <2μM, were also modestly higher within the thin layer (Fig. 7). In contrast, phosphate concentration increased sharply with depth beginning near the position of the thin layer where it was always > 5µM (Fig. 7).

The available temperature-salinity niche space expanded during this period, driven by warmer and saltier water above the pycnocline beginning in July (Fig 4D-E). However, the subsurface thin layer remained at temperatures similar to those observed during bloom development phase (14°C-18°C, Fig. 6). This general pattern was disrupted by three ephemeral mixing events on June 24, 27, and 29 (Fig. 3). The first two were associated with a shift in wind direction from southwesterly to northwesterly. The mixing event on the 29^th^ was associated with a slight cooling and freshening of the pond. This last mixing event coincided with cool overnight air temperatures (low of 14°C), but was not associated with any rainfall, changes in wind, or decreases in atmospheric barometric pressure (Fig. 3, Supporting Information Fig. 2). In each case, the dense thin layer re-established itself within less than one day. None of these changes in thin layer stability were associated with the spring-neap tidal time scale (Supporting Information Fig. 2A).

### Bloom decline following cessation of growth (July 5-12)

Standing stock began to decline when growth rates fell to near-zero, with vegetative and sexual reproduction contributing equally to that limited growth. Still, the *D. acuminata* thin layer remained cohesive (mean FWHM 67 m) and relatively deep (4–4.5 m depth) and was unaffected by tidal or diel cycles. Chemical measurements from July 9 similarly showed little change in the ammonium and pH profiles (10-27 µM NH_4_^+^ maximum and pH < 7.2 near the thin layer; Fig. 7).

Salt Pond’s surface water warmed through this period (19-28°C), and the whole pond became more saline (salinity 29.5-30.8). Still, temperatures within the thin layer remained similar to those observed during earlier phases (15-17°C; Fig. 6). On July 5, coincident with the spring tide, an intrusion of cold, salty water entered the pond, disrupting the thin layer (Fig. 3C-D). By the next available observations on July 8, the thin layer had re-formed. Physical conditions during this period were conducive to thin layer re-establishment and maintenance.

Specifically, tidal amplitude was low during the neap tide and winds were light and directionally consistent, driving relatively stable water column conditions and slowing the bloom’s decline (Fig. 3, Supporting Information Fig. 2).

The estimated residence time of *Dinophysis* cells during the period was ∼26 days (Supporting Information Fig. 4), substantially longer than residence times predicted for *Alexandrium* in Salt Pond if cells were well-mixed (∼2 days) or vertically migrating between 2.5 – 5m depth with weak hydrodynamic forcing (∼10 days) (Ralston et al. 2015). The extended residence time is consistent with the concentration *of D. acuminata* at depth, where cells experience reduced tidal exchange.

### Rapid termination (July 13-23)

Beginning July 13, standing stock declined rapidly. Intrinsic growth continued to be near-zero with no sexual reproduction observed. The *D. acuminata* thin layer was still cohesive but slightly more dispersed (FWHM 75cm) and shoaled into slightly warmer water (∼18-20°C) around 3.5m deep (Fig. 3B-C, Fig. 6). This change in the thin layer structure did not coincide with any notable shift in wind or tidal dynamics.

The structure of the ammonium profile changed substantially by bloom’s end (July 23). Ammonium concentrations increased monotonically with depth instead of peaking locally at the thin layer. However, concentrations remained quite high throughout the profile (∼8-12µM; Fig. 7). Likewise, pH values increased to >7.5 through the pond’s full depth (Fig. 7).

The residence time estimated from the best decay fit throughout this period was ∼5.6 days, lower than during the decline phase but still longer than expected from a well-mixed population (Ralston et al. 2015). Because this estimate also includes biological losses (e.g., grazing, infection, or senescence), it represents a conservative estimate of physical residence time. Image pixel grayscale intensity did not change and per-cell fluorescence was similar to the level recorded during the bloom’s initial development (Supporting Information Fig. 1A-B) and there was little other evidence of widespread biological losses in the IFCB image record. Therefore, shoaling of the thin layer appeared to drive standing stock decline by increasing advective losses during tidal exchanges. Grazing by larger micro- and mesozooplankton may have also contributed to losses but these were not observable by IFCB, putting these higher trophic level dynamics outside the scope of this study.

## Discussion

This study documented an intense, long-lasting *Dinophysis* bloom that persisted as a subsurface thin layer in a coastal kettle pond. The bloom duration was similar to that of previous blooms observed in this site, but recorded cell densities were higher (Ayache et al., 2023; Sung-Clarke et al., 2026). The bloom’s dynamics were observed through coupled continuous profiling and in-situ imaging and were well-resolved due to *Dinophysis*’ exceptional density, dominance in the plankton community, and phycoerythrin-rich kleptoplastids, which enabled vertical tracking. Because *Mesodinium* were scarce in the pond, *Dinophysis* could not renew their kleptoplasts through ingestion of new prey; hence, photosynthetic competence was observed declining throughout the course of the bloom. The amplitude of the cells’ vertical migrations declined with decreasing photosynthesis, indicated by reduced oxygen production and increased DIC concentrations within the thin layer, ultimately ceasing migration altogether. Cells settled in a sub-pycnocline thin layer that lay deeper than the tidal channel connecting the pond to the greater Nauset Marsh and Atlantic Ocean, driving their selective retention in the pond. This retention enabled observation of how prey deprivation altered cell metabolism and behavior alongside evolving thin layer chemistry. As the water occupied by *Dinophysis* acidified and was enriched in inorganic nitrogen and carbon, cells persisted in that thin layer, revealing an affinity for eutrophic conditions.

### Trophic mode linked to vertical swimming behavior

The coincidence in timing between the cessation of photosynthesis and vertical migration suggests a link between *D. acuminata* metabolic state and vertical migration behavior: cells migrated vertically while photosynthetically active, then maintained a static thin layer depth once photosynthesis declined. Over the past few decades, observations of vertical migration in *Dinophysis* have been inconsistent. For instance, in the Galician Rías, Spain, *Dinophysis* cells have been observed making diel vertical migrations of up to 7 m (Villarino et al. 1995; Reguera et al. 2003) or holding a stable depth position in different years (Pizarro et al. 2008; González-Gil et al. 2010). Similarly, non-migrating *Dinophysis* thin layers have been reported in the Baltic Sea and Patagonian fjords (Sjöqvist and Lindholm 2011; Baldrich et al. 2021). These conflicting reports may reflect differences in population metabolic state. Recently fed, photosynthetically active cells may benefit from vertical migrations that optimize light exposure and other ecological needs, whereas static thin layer formation may be more advantageous as cells lose photosynthetic capability.

Prior studies of vertical swimming and thin layer formation in dinoflagellates have focused primarily on phototrophic bloom-forming species that exploit opposing vertical gradients of light and nutrient availability (MacIntyre et al. 1997; Ault 2000; Zheng et al. 2023). Mixotrophic species like *Dinophysis* must additionally optimize prey encounter, prey capture, and predator avoidance (Hays et al. 2001; Sheng et al. 2007). In *Dinophysis*, these behavioral trade-offs are further complicated by its prey specialization. Until *Dinophysis* consumes *Mesodinium* and harvests its plastids, its behavior is driven by heterotrophic demands. Once kleptoplasts are acquired, behavior is also shaped by phototrophy. In this study, *Dinophysis* exhibited this apparent shift: cells produced oxygen and migrated vertically when photosynthetically active but later remained in a sub-pycnocline thin layer as photosynthetic activity declined (Fig. 8).

**Figure 8.**
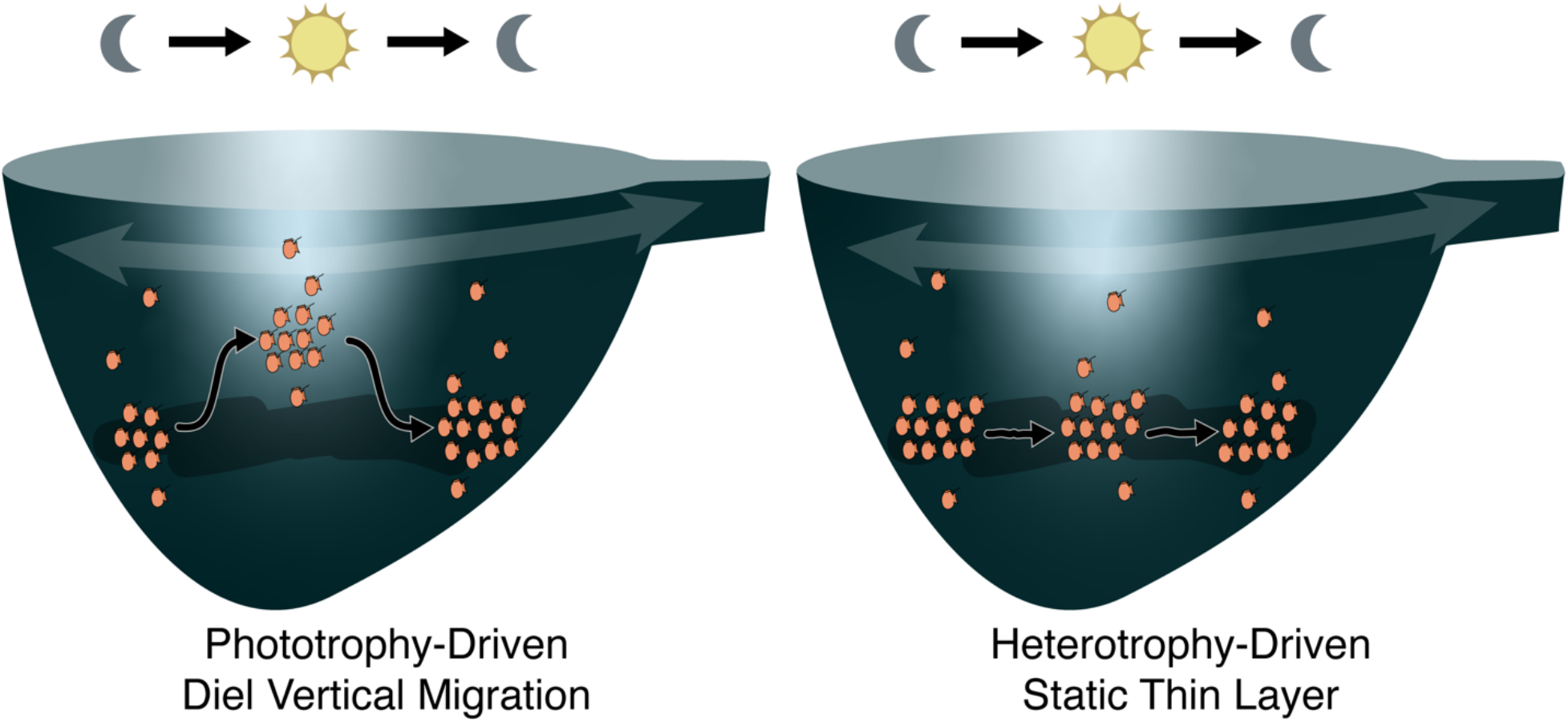
Conceptual model of the interaction between growth, vertical migration, and bloom retention for *Dinophysis acuminata* during the 2024 Salt Pond bloom. During the growth phase, cell behavior is driven by phototrophy: migrating upward during the day to access more light and down at night to access deeper nutrient reservoirs. During the peak of the bloom, cells form a static thin layer associated with a high ammonium maximum and low pH but are no longer vertically migrating. This thin layer depth keeps cells largely outside the tidally exchanged surface water mass, promoting their retention in the tidal kettle pond.

Relative to migrations reported in the Galician Rías, vertical migration ambits were smaller (∼1m versus 5–7 m; Villarino et al. 1995), likely reflecting greater light attenuation in Salt Pond. This short migration distance still represented a large change in light availability and therefore potential carbon acquisition. At night, cells swam downward to below the pycnocline where ammonium and dissolved organic nitrogen are typically elevated (Crespo et al. 2011), better supporting their growth (Hattenrath-Lehmann and Gobler 2015; García-Portela et al. 2020). In contrast, as photosynthetic activity rapidly declined among cells in Salt Pond, *Dinophysis* ceased their upward migrations during daytime. Notably, this behavioral shift was not accompanied by obvious morphological degradation: neither image pixel gray level nor the phycoerythrin-per-cell ratio changed (Supporting Information Fig. 1), suggesting that plastids were retained but no longer functioned.

Thin layer depth was not consistently associated with a specific light level, temperature or salinity (Fig. 3C-D, Fig. 5A). Rather, cells held a position just below the pycnocline, above the anoxic layer, and coincident with a local ammonium maximum. This position below the pycnocline constrasts with observations from other systems, where *Dinophysis* thin layers were recorded at or above it (Alves and Mafra 2018; Díaz et al. 2021), suggesting that cells also do not aggregate at a fixed position relative to the pycnocline. Nevertheless, the frequent association of *Dinophysis* thin layers with pycnoclines across multiple systems underscores the importance of water-column stratification for thin layer formation and persistence.

The most likely driver of stationary thin layer formation was loss of photosynthetic competency and reversion to dependence on heterotrophy. Thin layer formation likely enhances *Dinophysis*’ ability to capture prey and reestablish photosynthetic competency. Because *Mesodinium* vertically migrate in response to light and time of day (Fenchel and Hansen 2006), *Dinophysis* may maintain vertical position and wait for *Mesodinium* to migrate through their thin layer, an “angler’s strategy” that has been supported by observations of migrating *Mesodinium* coinciding with non-migrating *Dinophysis* in the Baltic Sea, Galician Rías, and Chilean fjords (Sjöqvist and Lindholm 2011; Díaz et al. 2019, 2025; Baldrich et al. 2021). A fixed vertical position may also facilitate the cooperative production and maintenance of mucus traps that *Dinophysis* use to ensnare their prey (Giménez Papiol et al. 2016; Jiang et al. 2018), and concentrate toxins that may facilitate prey capture (Mafra et al. 2016). This passive ambush strategy is energetically efficient between rare feeding opportunities and likely well suited to capture of fast-swimming *Mesodinium* prey (Jiang et al. 2018). Consequently, *Dinophysis* cells may aggregate as tightly as physical conditions allow to maximize prey encounter and capture. Although *Mesodinium* were absent from Salt Pond during this study, maintenance of a dense thin layer could maximize capture success when ephemeral patches of migrating prey are encountered.

### Metabolism shapes thin layer chemistry

As the *Dinophysis* bloom shifted away from phototrophy into its heterotrophy-driven persistence phase, the thin layer became characterized by persistently low pH and locally elevated ammonium and DIC. Community respiration by *Dinophysis* – the dominant microplankton (Fig. 2A) – and their associated microbial community depleted oxygen, increased dissolved inorganic carbon, and remineralized organic nitrogen into ammonium. At the same time, the loss of photosynthetic competency diminished overall demand on the ammonium pool (García-Portela et al. 2020). Nitrification from ammonium into nitrite is inhibited by hypoxia, and is often relatively slow compared to ammonification of organic nitrogen in eutrophic coastal environments (Herbert 1999). Inhibition of nitrification during this bloom was supported by the relatively small increase in nitrate and nitrite despite persistently high ammonium (Fig. 7). As the bloom terminated, the thin layer ammonium, pH, and DIC structure dissipated, further supporting a critical role for *Dinophysis* in driving changes to local thin layer chemistry.

The thin layer chemistry observed here contrasts with typical algal bloom chemistry. Phototrophic bloom-formers deplete inorganic nutrients, increase dissolved oxygen, and deplete dissolved inorganic carbon, raising pH (Cloern 1996). On the other hand, *Dinophysis*—through loss of photosynthetic competency—became associated with increased inorganic nutrients and carbon, decreased dissolved oxygen, and acidification. These observations are consistent with recent observations of dense *D. acuminata* blooms in two New York estuaries amid intensifying nocturnal hypoxia and acidificiation (Farrell et al. 2026). For phototrophic species, those conditions typically only develop after bloom collapse and decay. The development and persistence of that niche concurrent with the *Dinophysis* bloom highlights that trophic mode of bloom-formers can fundamentally affect local chemical environments.

Despite acidic conditions, the *Dinophysis* thin layer persisted. Mixing events temporarily disrupted it several times (Fig. 4), but the cells repeatedly re-formed the thin layer in subsurface acidified waters, indicating tolerance or even a preference for that niche. This acidified, nutrient-rich niche may in fact generally support *Dinophysis* persistence. Because *Mesodinium* readily assimilates dissolved organic and inorganic nutrient sources (Wilkerson and Grunseich 1990) and are highly tolerant of low pH environments (Eriksen et al. 2023), the microenvironment the *Dinophysis* layer creates in its heterotrophy-limited state in may help lure its prey while at the same time reducing its own exposure to predation, e.g. from less acid-tolerant micro- or mesozooplankton (Wyeth et al. 2022). As a result, continued vertical positioning in the nutrient-rich, acidified thin layer likely reflects optimization of prey capture while offering potential secondary physiological and ecological benefits.

Coastal eutrophication drives high nutrients, acidification, and hypoxia (Howarth et al. 2011), the same conditions *Dinophysis* engendered and persisted through in Salt Pond. Their demonstrated persistence throughout such conditions during the bloom suggests a potential competitive advantage in eutrophic environments, enhancing risk of *Dinophysis* blooms and DSP.

### Consequences of vertical distribution on coastal blooms

The sub-pycnocline, sub-inlet vertical position of *Dinophysis* cells insulated them from tidal dilution. Despite the pond’s volumetric residence time of ∼ 2 days (Anderson and Stolzenbach 1985), tidal exchange primarily affects surface waters, extending the effective residence time for *D. acuminata* during this bloom to several weeks even after growth stopped. This is a conservative estimate, as it assumes all losses during bloom decline were via export from the pond. Previous models of *Alexandrium catenella* blooms in Salt Pond showed that diel vertical migration increased its residence time up to about 10 days, as downward migration at night similarly insulated it from tidal exchange (Ralston et al. 2015). Unlike *A. catenella*, which has its own plastids and is obligately phototrophic, prey-starved *Dinophysis* do not migrate upward during the day, even further insulating it from tidal exchange and exacerbating its trapping within Salt Pond. When the thin layer shoaled into shallower, more readily exchanged water, bloom termination occurred quickly, supporting physical export as the dominant loss mechanism, rather than as a direct consequence of physiological decline associated with starvation. Though cell morphology did not noticeably change, the thin layer shallowing still reflected a change in cell behavior, since there was no evidence of increased physical mixing or relaxation of stratification at termination onset.

These observations highlight how vertical swimming can interact with stratification and coastal bathymetry to generate intense blooms. They are consistent with previous studies demonstrating that stratified environments enhance *Dinophysis* retention and bloom development by promoting cell aggregation and limiting dispersal (Díaz et al. 2013, 2021). Salt Pond is a surprising location for such dense *Dinophysis* blooms given the rarity of their obligate prey; yet *Dinophysis* swimming behavior and stable stratification effectively concentrate these cells in this relatively deep terminal kettle hole. As many *Dinophysis* blooms have been observed without prey, physical trapping mechanisms such as this help explain frequent observations of mismatch between predator and prey occurrence.

Projected increases in stratification in the region are likely to increase the frequency and duration of *Dinophysis* blooms (Boivin-Rioux et al. 2022). Longer bloom persistence and toxin exposure could exacerbate the ecological, public health, and socioeconomic impacts of *Dinophysis* in the region (Pease et al. 2022; Ayache et al. 2025). The vertical distribution of *Dinophysis* also has consequences for bloom detection and monitoring programs. When *Dinophysis* do not migrate to the surface during the day, as many phototrophic dinoflagellates do (Brosnahan et al. 2017; Zheng et al. 2023), they evade detection from most HAB monitoring programs that target surface waters.

### Development of prey-free Dinophysis blooms

While coastal trapping explains the intensification and persistence of this bloom, it does not explain its initiation. The common predator-prey mismatch observed in *Dinophysis* blooms has led to some speculation that *Dinophysis* can rely on alternative prey (Reguera et al. 2012; Díaz et al. 2025), but the rapid decline in growth throughout the 2024 Salt Pond bloom suggests that the absence of *Mesodinium* constrained *Dinophysis* growth. Instead, the initial high growth rates indicate that the *Dinophysis* cells that seeded this bloom were well-fed. As *Mesodinium* were not observed within the pond, the bloom was likely not initiated locally, but instead tidally imported from outside Salt Pond. This scenario is consistent with the “pelagic seed bank” hypothesis (Smayda 2002) in which coastal blooms of holoplankton are initiated by offshore populations entrained in frontal zones, and has been hypothesized as a source of *Dinophysis* blooms in the Gulf of Mexico (Harred and Campbell 2014). *Mesodinium* have been observed most frequently in the Gulf of Maine from the fall through spring (Sanders 1995; Brownlee 2017); these populations may sustain coastal *Dinophysis* populations. Though no *Dinophysis* population has been observed offshore near this site, identifying such a source, as well as any offshore *Mesodinium* prey population that they feed on, would improve understanding and prediction of coastal *Dinophysis* blooms.

## Acknowledgements

We would like to thank the Cape Cod National Seashore for allowing us to conduct this study in Salt Pond. We are also grateful to Mrunmayee Pathare, Michael Staiger, Ryan Govostes, and Nathan Figueroa with their assistance with the deployment of the PhytO-ARM observing platform in Salt Pond. Claude Haiku (version 4.5) was used during revision to streamline and improve clarity of text. This paper is a result of research funded by the National Oceanic and Atmospheric Administration (NOAA) National Centers for Coastal Ocean Science Competitive Research Program under award NA19NOS4780182 to the Virginia Institute of Marine Science, the NOAA Ocean Acidification Program under award number NA22NOS4780170, and the Woods Hole Center for Oceans and Human Health (National Science Foundation grant OCE-1840381 and National Institutes of Health grant NIEHS-1P01-ES028938-01 to the Woods Hole Oceanographic Institution). Contributions by W.G.Z. were funded by the Qiushi Feiying project at Zhejiang University. This is ECOHAB publication number 1153.

