## Supporting Information for "Cessation of diel vertical migration by an inshore dinoflagellate bloom under prey deprivation"

Serena Sung-Clarke[1,2], Nour Ayache[1,3], Wenguang Zhang[4], David K. Ralston[1], Evan Lechner[1], Zhaohui Aleck Wang[1], Juliette L. Smith[5], Collin Roesler[6], Susan Drapeau[6], Mengmeng Tong[4], Michael Brosnahan[1]

[1] Woods Hole Oceanographic Institution, Woods Hole, MA, USA, [2]MIT-WHOI Joint Program in Oceanography/Applied Ocean Science & Engineering, Cambridge and Woods Hole, MA, USA, [3] French Research Institute for Exploitation of the Sea (Ifremer), Nouméa, New Caledonia, France, [4] Ocean College, Zhejiang University, Zhoushan, Zhejiang, China, [5] Virginia Institute of Marine Science, William & Mary, Gloucester Point, VA, USA, [6] Bowdoin College, Brunswick, ME, USA

**Supporting Information Text S1. DIC and pH measurement details.**

Discrete DIC sample analyses were conducted via a DIC auto-analyzer (AS-C6L DIC multi-sample analyzer, Apollo SciTech Inc., Delaware, USA). For this method, each sample was acidified with 3% phosphoric acid and then purged of total $CO_2$ by ultra-pure nitrogen gas. DIC was then measured by infrared $CO_2$ analyses of the purged $CO_2$ utilizing a LI-7815 $CO_2$/$H_2O$ Trace Gas Analyzer (LI-COR Environmental, Nebraska, USA). Instruments were calibrated using a secondary reference standard certified by Certified Reference Materials (CRM) provided by Dr. A. Dickson at Scripps Institute of Oceanography. The precision and accuracy for DIC analyses were ±0.1%.

Discrete pH sample analyses were conducted with an Agilent 8453 following standard spectrophotometric procedures (Dickson et al. 2007; Douglas and Byrne 2017). Each sample was added into a pair of two 10 cm long cylindrical cells via a 60 mL syringe. Each cell was rinsed three times with 5 mL of sample, and the remaining 45 mL was used to fill and then overflow the cell. Cells were then held in a temperature-controlled manifold for a minimum of one hour to allow for temperature equilibration at 25.0 ±0.1 °C. For each cell, reference absorbances were recorded before the addition of 20 µL of 4 mM purified m-cresol purple (mCP). Absorbances were then measured at three wavelengths (434 nm, 578 nm, and 730 nm). The ratio between absorbances at 578 nm and 434 nm was determined and corrected for any baseline drift recorded at 730 nm. The corrected absorbance ratio was then used to calculate sample pH on the total scale using the mCP model as described by Douglas and Byrne (2017). Overall pH measurement uncertainty was estimated as described by Song et al. (2020) with a mean uncertainty of ±0.006, based on duplicate sample analyses (n = 63).

**Supporting Information Text S2. Dissolved nutrient sample analysis.**

Dissolved nutrient samples were slowly thawed overnight in a refrigerator the day before analysis. Reagents and standards were made fresh for each set of analyses. Ammonium concentrations were determined following Standard Methods 4500-NH3-G (19$^{th}$, 20$^{th}$, and 21$^{st}$ Edition) and 4500-NH3-H (18$^{th}$ Edition). Combined nitrate and nitrite, or total oxidized nitrogen $NO_X$, concentrations were determined from the EPA Method 353.2 Revision 2.0 (1993) and Standard Methods 4500 NO3F, 18$^{th}$ and 19$^{th}$ Editions. Phosphate concentrations were determined following EPA Method 365.1, Revision 2.0 (1993). Silicate concentrations were determined using the method based on USGS I-2700-85 and Standard Methods 4500-Si-F 18$^{th}$ and 19$^{th}$ Editions.

**Supporting Information Text S3. Estimating role of export in bloom decline.**

Estimates of residence time, $\tau$, were calculated by fitting exponential decay function to standing stock estimates, $N$, throughout the bloom.

$$N(t) = N_0 e^{-\frac{t}{\tau}} \qquad (7)$$

Residence times were estimated over two periods of time when growth rates were near-negligible (<0.05 day$^{-1}$): when thin layer was maintained and standing stock decrease was modest, and upon a sharp decline in population size. This is a lower-bound estimate of residence time because it assumes loss was solely attributed to export. Other factors that may have driven cell loss—e.g., grazing, infection, senescence—are neglected. These residence time estimates were compared to residence times calculated from hydrodynamic model simulations of phytoplankton in Salt Pond with varying swimming behavior and forcing conditions (Ralston et al. 2015).

**Supporting Information Text S4. Whole cell counts from within and outside Salt Pond.**

To assess tidal exchange of *Dinophysis* cells, water samples were collected from both the observing platform inside the pond and outside of Salt Pond in Nauset Marsh (Fig. 1). On June 7 and 12, triplicate 2-L bottles were taken from the surface outside Salt Pond during flood tide and from the phycoerythrin maximum inside Salt Pond during slack high tide. On five sampling trips from June 20 - 23, triplicate 2-L bottles were collected from the surface outside Salt Pond during both flood and ebb tides, along with 2-L bottle samples from inside Salt Pond at the surface and phycoerythrin maximum. Samples were concentrated over 15µm Nitex mesh to 15mL volume, fixed with Acid Lugol's Solution, and transported back to the lab, where they were counted on a 1-mL Sedgewick Rafter.

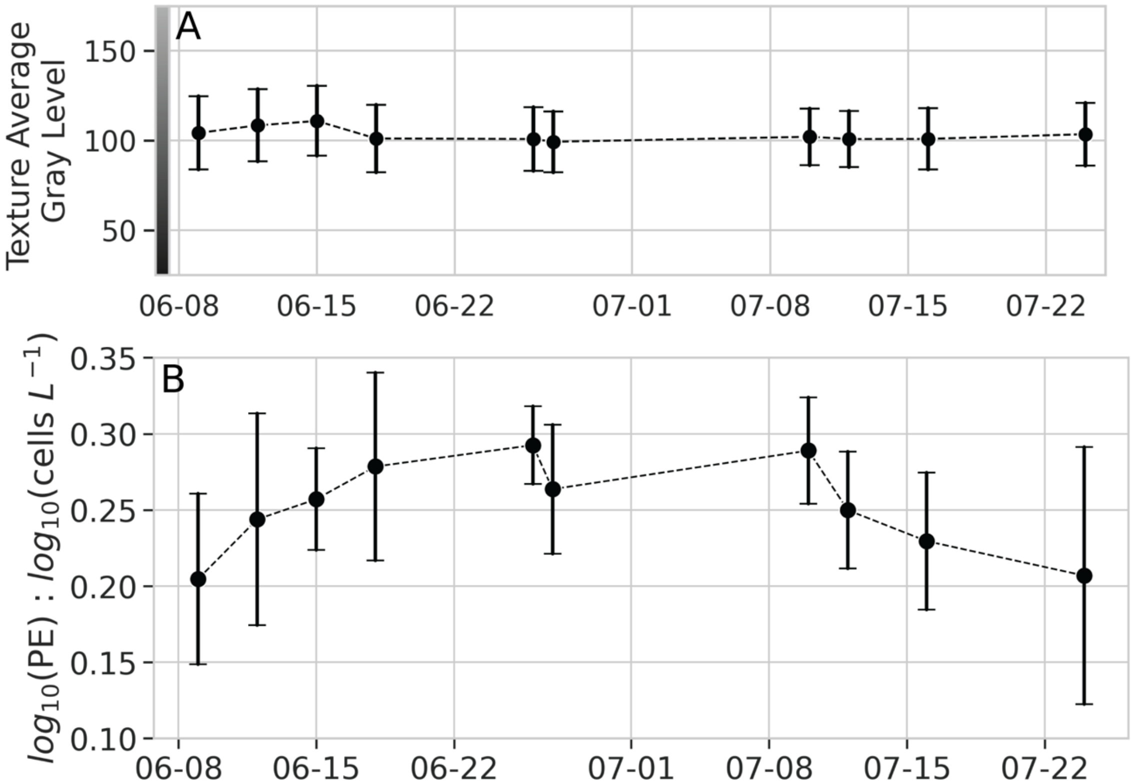


**Supporting Information Figure 1. (A)** The distribution of texture average gray level (TAGL) across fully annotated days in the bloom. The colorbar on the left represents the grayscale associated with each TAGL value. **(B)** Mean and standard deviation of the ratio between log-transformed phycoerythrin and cell concentration.

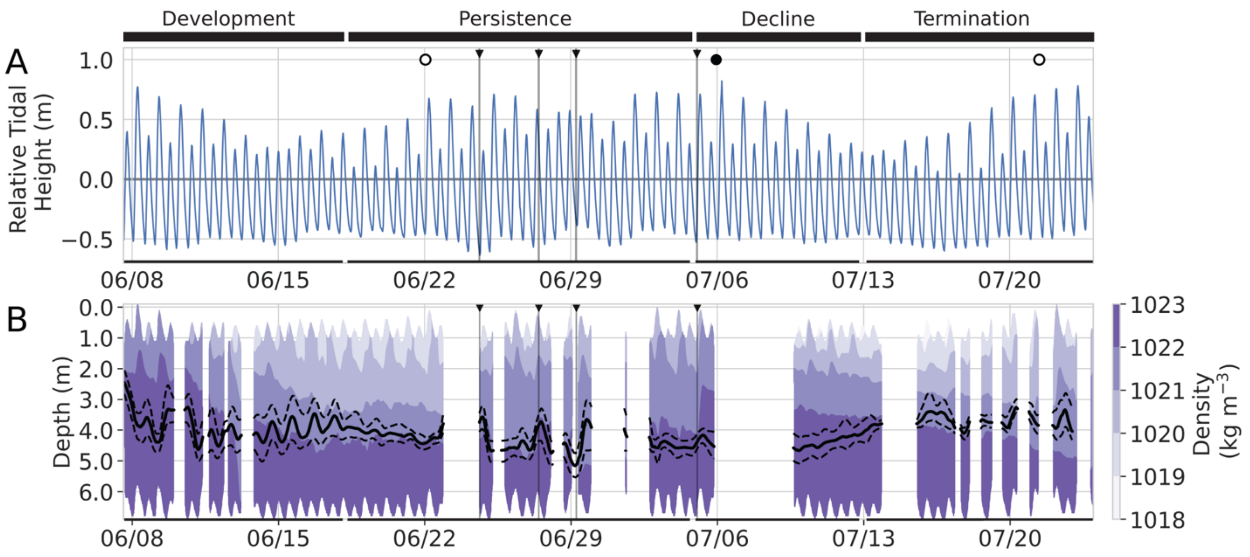


**Supporting Information Figure 2. (A)** Tidal height relative to the mean in the logger record during the 2024 *Dinophysis* bloom. Open and closed circles represent date of the full and new moon, respectively. **(B)** Density profile throughout the bloom, with the black solid and dashed lines representing the smoothed depth of the phycoerythrin maximum and its upper and lower depth bounds. Depth is the depth below the mean sea level surface height. Vertical lines highlight four thin layer disruptions during the peak of the bloom.

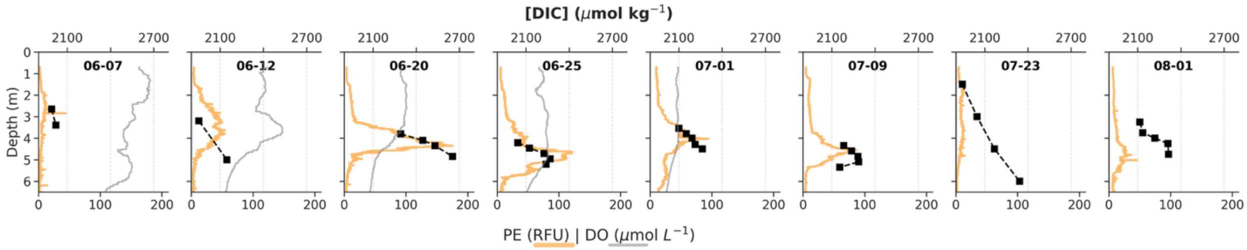


**Supporting Information Figure 3.** Dissolved inorganic carbon (DIC) profiles across 8 days during the 2024 Salt Pond bloom compared to the profiles of phycoerythrin and dissolved oxygen, where available.

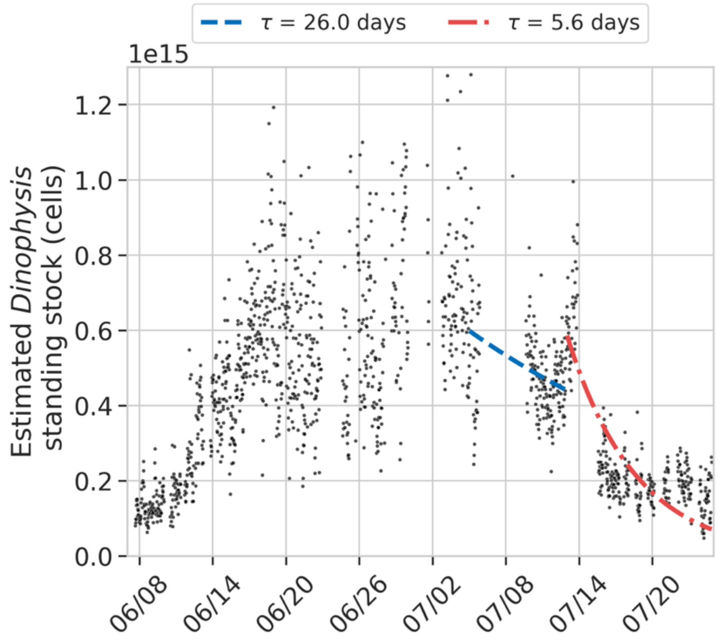


**Supporting Information Figure 4.** Salt Pond standing stock estimates (points) from the beginning of the bloom's decline until the end of the observation period. Dashed lines represent the best exponential decay fit for the decline phase and termination phases of the bloom.

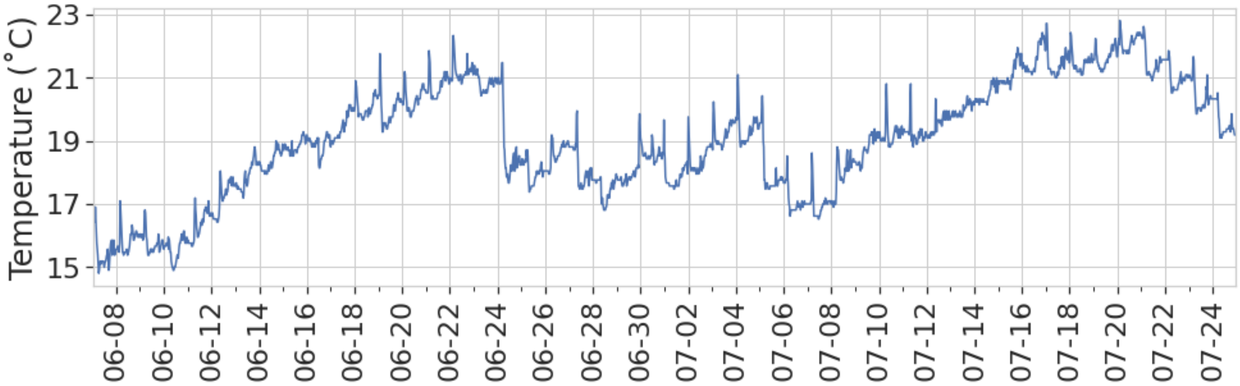


**Supporting Information Figure 5.** Temperature recorded by a HOBO logger that was moored to the bottom of Salt Pond near the inlet (~2m depth).

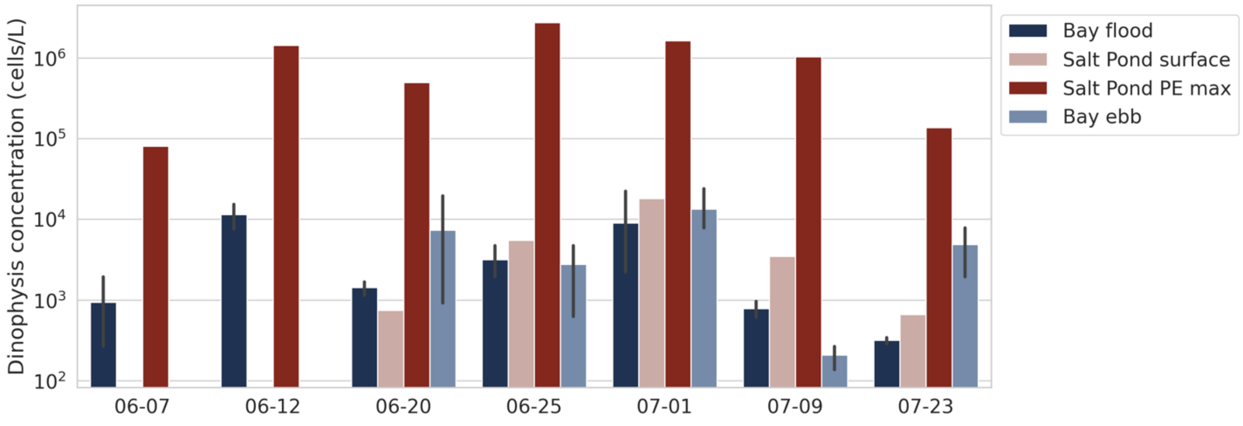


**Supporting Information Figure 6.** *Dinophysis* concentrations based on cell counts from the surface and phycoerythrin maxima in Salt Pond, as well as from Salt Pond Bay during flood and subsequent ebb tide.

**Supporting Information Citations**

Clayton, T. D., and R. H. Byrne. 1993. Spectrophotometric seawater pH measurements - Total hydrogen ion concentration scale calibration of m-cresol purple and at-sea results. Deep-Sea Res. (I) 40: 2115-2129.

Dickson, A. G., C. L. Sabine, and J. R. Christian. 2007. Guide to best practices for ocean CO2 measurements. PICES Special Publication.

Douglas, N. K., and R. H. Byrne. 2017. Spectrophotometric pH measurements from river to sea: Calibration of mCP for 0 <= S <= 40 and 278.15 <= T <= 308.15 K. Mar. Chem. 197: 64-69

Ralston, D. K., M. L. Brosnahan, S. E. Fox, K. D. Lee, and D. M. Anderson. 2015. Temperature and Residence Time Controls on an Estuarine Harmful Algal Bloom: Modeling Hydrodynamics and Alexandrium fundyense in Nauset Estuary. Estuaries Coasts **38**: 2240–2258. doi:10.1007/s12237-015-9949-z

Song, S., Z. A. Wang, M. E. Gonneea, K. D. Kroeger, S. N. Chu, D. Li, and H. Liang. 2020. An important biogeochemical link between organic and inorganic carbon cycling: Effects of organic alkalinity on carbonate chemistry in coastal waters influenced by intertidal salt marshes. Geochim. Cosmochim. Acta 275: 123-139.
